# The neural encoding of information prediction errors during non-instrumental information seeking

**DOI:** 10.1101/179507

**Authors:** Maja Brydevall, Daniel Bennett, Carsten Murawski, Stefan Bode

## Abstract

In a dynamic world, accurate beliefs about the environment are vital for survival, and individuals should therefore regularly seek out new information with which to update their beliefs. This aspect of behaviour is not well captured by standard theories of decision making, and the neural mechanisms of information seeking remain unclear. One recent theory posits that valuation of information results from representation of informative stimuli within canonical neural reward-processing circuits, even if that information lacks instrumental use. We investigated this question by recording EEG from twenty-three human participants performing a non-instrumental information-seeking task. In this task, participants could pay a monetary cost to receive advance information about the likelihood of receiving reward in a lottery at the end of each trial. Behavioural results showed that participants were willing to incur considerable monetary costs to acquire early but non-instrumental information. Analysis of the event-related potential elicited by informative cues revealed that the feedback-related negativity independently encoded both an information prediction error and a reward prediction error. These findings are consistent with the hypothesis that information seeking results from processing of information within neural reward circuits, and suggests that information may represent a distinct dimension of valuation in decision making under uncertainty.

## Introduction

Seeking information is an important drive of behaviour, and a key component of effective decision making under uncertainty^1^. However, normative decision theory, which assumes that the value of information resides in its instrumental utility for acquiring future rewards^2–4^, provides a poor description of information seeking in humans and other animals. In particular, such theories cannot account for findings showing that animals place a positive value on information that resolves uncertainty but which cannot be used to affect future tangible outcomes (termed non-instrumental information). Human participants, for instance, display a clear preference for acquiring non-instrumental information about both aversive and appetitive future events^5–7^ and many species, including humans, exhibit a willingness to sacrifice part of an uncertain future reward in exchange for non-instrumental information about the reward’s likelihood^6, 8–10^. These behavioural findings indicate that animals treat information as though it were of intrinsic value (cf. Grant, Kajii and Polak, 1998^11^).

One recent proposal, the ‘common currency’ hypothesis, is that the intrinsic value of information might result from common neural substrates for processing of rewarding and informative stimuli^12^. Neural recordings from non-human primates have demonstrated that non-instrumental information is encoded within brain regions typically associated with reward processing, such as the dopaminergic midbrain^13^, lateral habenula^12^ and orbitofrontal cortex^9^. Notably, Bromberg-Martin and Hikosaka (2011) reported that in response to informative stimuli, neurons in macaque lateral habenula encoded both reward prediction errors (RPEs; the signed difference between expected and actual reward) and information prediction errors (IPEs; the signed difference between expected and actual information). Similarly, functional magnetic resonance imaging (fMRI) in humans has revealed that the delivery of information is associated with increased blood-oxygen-level dependent signals within brain regions typically associated with reward processing, such as the striatum^14, 15^. This resemblance suggests a common neural coding scheme for information and primary reward, which might result from mechanisms such as an intrinsic reward value of information^12^ or boosting of anticipatory utility by reward prediction errors associated with informative stimuli^16^.

To date, many predictions of the common currency hypothesis of information valuation have not been investigated in humans. To address this question, the present study recorded the electroencephalogram (EEG) from human participants completing a non-instrumental information seeking task, assessing willingness-to-pay for non-instrumental information^6^. On each trial, a lottery was drawn in which participants had an equal probability of winning (receiving 20 cents) or losing (receiving 0 cents). Prior to the lottery draw, participants could choose to view either an informative stimulus, which imparted early information about the lottery outcome, or a non-informative stimulus, which was perceptually identical to the informative stimulus but imparted no information about the lottery outcome. To assess participants’ willingness to pay for non-instrumental information, a variable cost was associated with viewing the informative stimulus, to be deducted from participants’ winnings in the case of a win outcome only. We then investigated whether IPEs associated with non-instrumental information were encoded in the feedback-related negativity (FRN) component of the event-related potential (ERP). According to one prominent theory, FRN amplitude reflects RPEs following the disinhibition of neurons in anterior cingulate cortex by mesencephalic dopamine neurons^17^. In support of this contention, it has been shown that FRN amplitudes are greater following negative RPEs than positive RPEs^18, 19^. Indeed, it has been proposed that the FRN could be reconceptualised as a ‘reward positivity’^20^ encoding the hedonic value of stimuli relative to expectations. Premised upon this dopaminergic RPE model of the FRN, the common currency hypothesis of information valuation therefore predicts that IPEs should also be encoded in the FRN, in a comparable fashion to RPEs.

## Results

### Behavioural results

One participant failed an attention check and was therefore excluded from further analysis (see Methods). Behavioural results (Fig. 1) replicated the overall findings of Bennett et al. (2016)^6^. A repeated-measures analysis of variance (ANOVA) revealed that preference for information was modulated by the cost of information (*F*(1.76, 36.99) = 58.02, *p <*.001,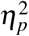 = 0.42). Participants displayed a strong preference for the informative stimulus when it was available at no cost (*t*-test against 0.5: *t*(21) = 16.96, *p <*.001), and a non-negligible preference for this stimulus when it was available at a cost (*t*-test against zero: *t*(21) = −3.55, *p* =.01). Preference for information decreased with increasing information cost (single-sample *t*-test of coefficients from a linear regression against zero: *t*(21) = −9.98, *p <*.001). There was also considerable inter-individual variability in preference for information as measured by overall proportion of choices to observe the informative stimulus (*M* =.39, range =.15-1). These results suggest that participants assigned an intrinsic value to the non-instrumental information imparted by the informative stimulus.

**Figure 1.**
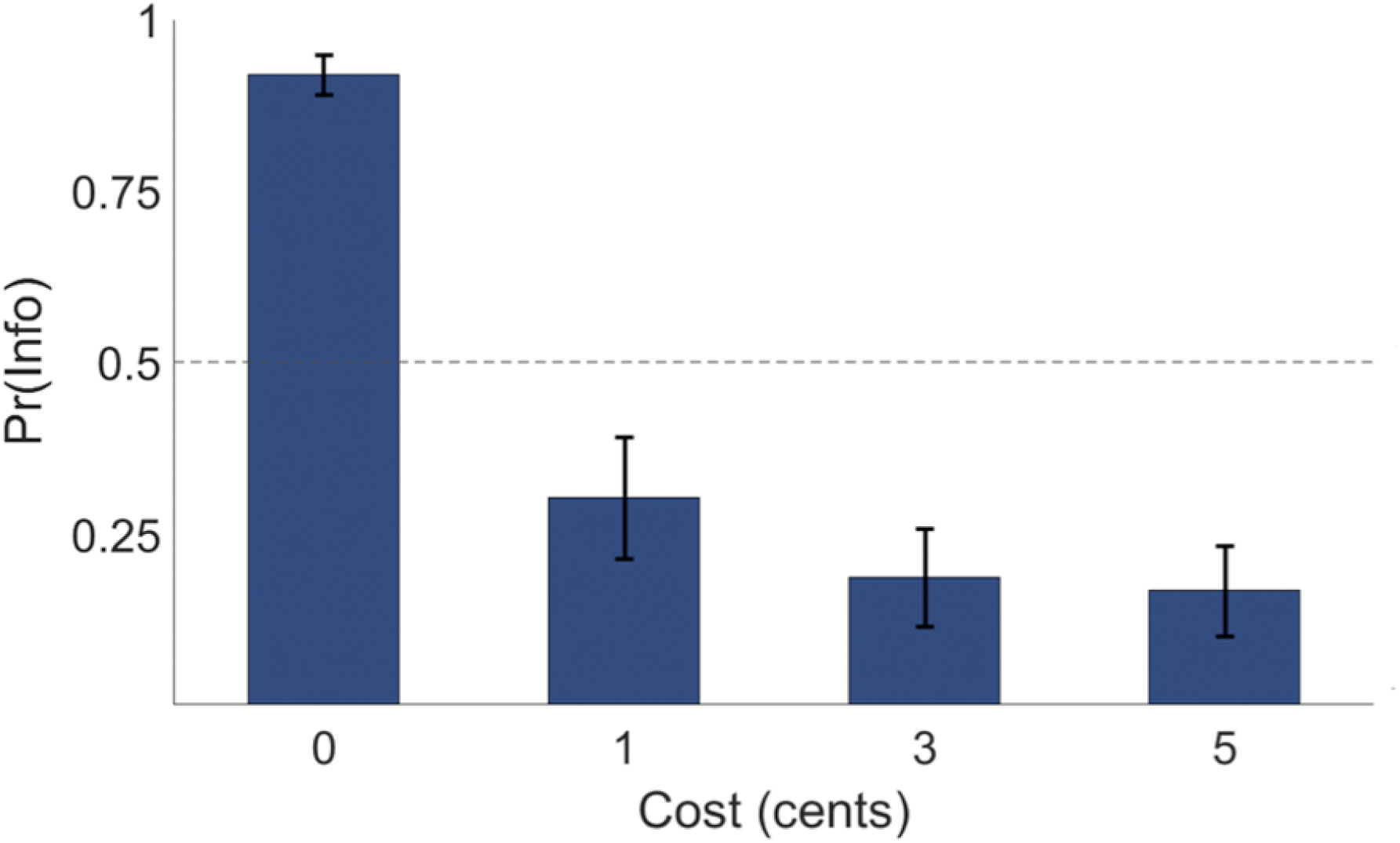
Behavioural results. Mean proportion of informative stimulus choices (denoted Pr(Info)) as a function of information cost. Error bars represent the standard error of the mean.

### ERP results

We investigated how RPEs and IPEs were encoded in the amplitude of the feedback-related negativity elicited by the presentation of informative cues (both card presentation and trial outcome screens; see Method for further information regarding trial structure). RPEs were calculated as the discrepancy between expected lottery winnings prior to observing the stimulus and actual expected lottery winnings after observing the stimulus:

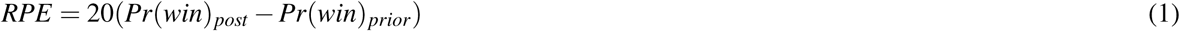

Similarly, IPEs were calculated as the difference between the actual information content of a stimulus *I* and its expected information content *I*_*expected*_ :

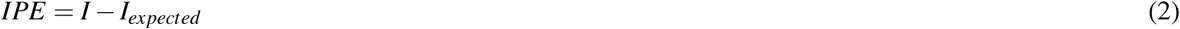

Information content was itself calculated as the reduction of belief entropy, as per information theory (see Method for further information regarding the computation of these variables). Analogous to RPEs, a positive IPE occurred upon presentation of stimuli that conveyed more information than expected, and vice versa for negative IPEs. Crucially, the equiprobability of red and black cards meant that IPEs and RPEs were statistically independent of one another by design.

#### Reward prediction errors

We first examined whether the amplitude of the FRN evoked by the presentation of informative cards encoded RPEs, as predicted by a prominent reinforcement learning theory^17^. In line with previous studies^19, 21^, we analysed FRN amplitudes using a 2 x 5 repeated-measures ANOVA with factors of RPE (positive, negative) and electrode (Fpz, AFz, Fz, FCz, Cz), which revealed a significant main effect of RPE on FRN amplitude (*F*(1, 14) = 6.09, *p* =.03, 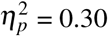), with negative RPEs associated with a more negative FRN amplitude compared to positive RPEs (see Fig. 2A). This indicates that the FRN encoded RPEs in a typical fashion in the present study.

**Figure 2.**
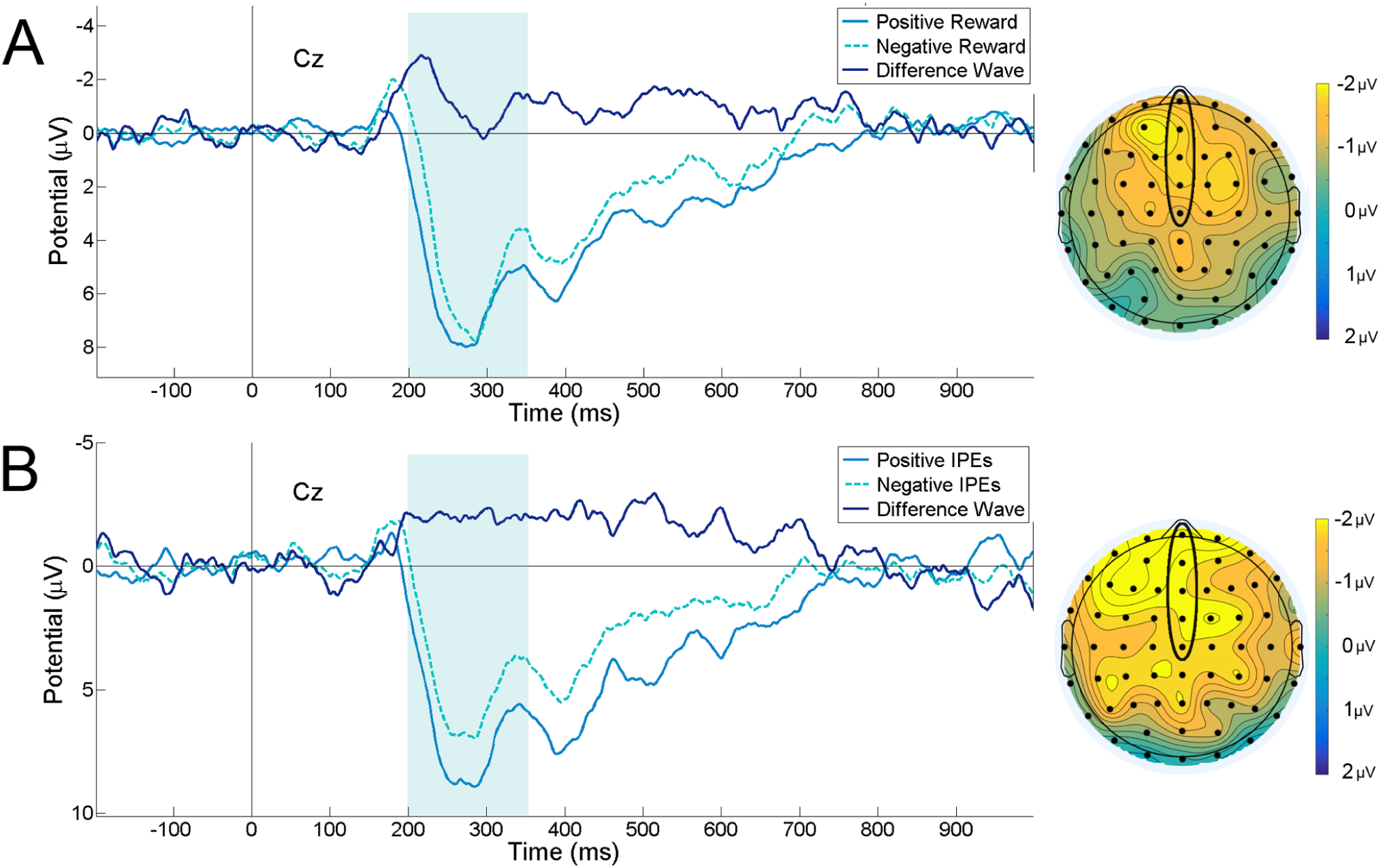
ERP waveforms and scalp maps. Grand average ERP waveforms and scalp maps for positive and negative reward prediction errors (A) and positive and negative information prediction errors (B) at electrode Cz. The dark blue waveform denotes a difference wave (negative - positive prediction errors), and the teal rectangle denotes the FRN measurement window (200–350ms). Scalp maps display the topography of the mean difference wave over this measurement window. Circled frontocentral electrodes are those at which FRN amplitude was calculated for analysis. In both grand average waveforms and scalp maps, negative voltages are plotted upwards.

We also observed a significant main effect of electrode on FRN amplitude (*F*(1.62, 22.61) = 53.66, *p <*.001,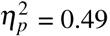); however, this effect did not interact significantly with the effect of RPE (*F*(1.83, 25.55) = 0.98, *p* =.43).

#### Information prediction errors

As with the RPE analysis, we used a 2 x 5 repeated-measures ANOVA to investigate the effects of IPE (positive, negative) and electrode (Fpz, AFz, Fz, FCz, Cz) on FRN amplitude. Analogous to the effect of RPE, we observed a significant main effect of IPE on FRN amplitude (*F*(1, 14) = 7.75, *p* =.01, 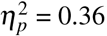), driven by significantly more negative FRN amplitudes in response to negative IPEs than to positive IPEs (see Fig. 2B). These results indicate that both IPEs and RPEs were encoded by the FRN: negative prediction errors—both RPEs and IPEs—both elicited more negative FRN amplitudes relative to positive prediction errors. As for the RPE analysis, we also found a significant main effect of electrode on FRN amplitude (*F*(1.71, 23.90) = 37.83, *p*= .001,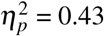), but no interaction between electrode and RPE (*F*(4, 56) = 0.19, *p* =.93).

To assess the generality of these findings, we next conducted an additional control analysis to determine whether a similar modulation of FRN amplitudes was observed when zero IPE events were also included in analysis. To this end, a 5 x 3 repeated-measures ANOVA was used to assess the within-participants effects of IPE (positive, negative, zero) and electrode on the amplitude of the FRN. As above, this new analysis revealed a main effect of IPE for the informative stimulus (*F*(2, 28) = 6.03, *p <*.01,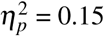). Consistent with the results of the main analysis, post-hoc paired-sample t-tests with Bonferroni correction for multiple comparisons indicated that this main effect was driven by a significantly more negative FRN for negative IPEs (*M* = 2.81, *SEM* = 0.66) than for positive IPEs (*M* = 4.94, *SEM* = 0.71; *p* =.04), as well as for negative IPEs relative to zero IPEs (*M* = 4.28, *SEM* = 0.59; *p* =.02).

#### Amount of information

We also examined whether the absolute amount of information delivered by stimuli was also encoded in the FRN. This involves looking at information independent of expectations; positive information can be defined as becoming more certain of the trial outcome (both more certain of winning and more certain of losing), whereas negative information involves becoming less certain of the trial outcome. It is important to note that amount of information will tend to be positively correlated with the sign of IPEs, since trials with a positive IPE are a subset of all trials with a positive amount of information, and vice versa for negative IPEs. As such, this analysis should be considered an incremental modification of the IPE analysis presented above, rather than a discrete inquiry.

Using a 2 x 5 repeated-measures ANOVA, we assessed the effect of information (positive, negative) and electrode (Fpz, AFz, Fz, FCz, Cz) on FRN amplitude. We found a significant main effect of information (*F*(1, 14) = 9.59, *p <*.01,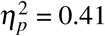), with more negative FRN amplitudes for negative information (greater uncertainty) relative to positive information (greater certainty). Again, we observed a significant main effect of electrode (F(1.69, 23.66) = 61.30, *p <*.001, 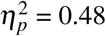), but no interaction effect between information and electrode (*F*(2.36, 32.97) = 0.87, *p* =.49). ERP waveforms for this analysis are presented in supplementary Figure S1.

#### Non-informative stimuli

As an additional control analysis, we next investigated whether the modulation of the FRN in response to RPEs and IPEs was unique to cards following a decision to view the informative stimulus, or whether similar patterns were observed for cards following a decision to observe the non-informative stimulus. To do this, we calculated RPEs and IPE variables for cards in each non-informative stimulus as though they had instead been an informative stimulus (because technically, according to the formulae set out in the Method section, IPEs and RPEs were always zero for cards in the non-informative stimulus).

We observed no modulation of the FRN by RPEs for the non-informative stimulus (*F*(1, 18) =.02, *p* =.90; see supplementary Figure S2). However, in an in interesting parallel to the effect of IPE in the informative stimulus, we also observed a small effect of IPE (*F*(1, 18) = 4.58, *p* =.046, 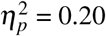; see supplementary Figure S3). There was no significant effect of absolute amount of information on FRN amplitude for non-informative stimuli (*F*(1, 18) = 0.47, *p* =.50; see supplementary Figure S4).

#### Outcome screen ERPs

There was no significant difference in FRN amplitudes between win and loss outcomes when analyses were conducted separately for outcome screens following an informative stimulus (*F*(1, 18) = 3.74, *p* =.07; see supplementary Figure S5) and for outcome screens following a non-informative stimulus (*F*(1, 17) = 0.11, *p* =.74; see supplementary Figure S6).

## Discussion

This study used an information seeking task to investigate human participants’ preference for non-instrumental information in decision making under uncertainty. Using EEG, we assessed how both reward prediction errors and information prediction errors were reflected in the feedback-related negativity component of the event-related potential. Behavioural results replicated the overall pattern of findings previously reported by Bennett and colleagues (2016), consistent with an intrinsic valuation of information (cf. Grant et al., 1998). That is, participants displayed a clear preference for acquiring non-instrumental information, despite the fact that this information was at times associated with a direct monetary cost. Analyses of the ERP evoked by informative stimuli revealed that RPEs and IPEs were both encoded in a comparable fashion in the amplitude of the FRN component.

ERP analyses showed that the modulation of the FRN during task events that elicited positive and negative IPEs was consistent with FRN modulation by positive versus negative RPEs. The FRN has traditionally been considered to encode correct and incorrect responses in tasks^17, 22^, as well as rewarding outcomes^21, 23^. As such, our ERP analyses show a striking parallel in FRN encoding of informative and rewarding outcomes. This is conceptually consistent with the finding that firing rates of single neurons in primates respond in the same manner to positive/negative IPEs as to positive/negative RPEs^12^. Since FRN amplitude is thought to be related to dopaminergic projections to the anterior cingulate cortex^24^, the modulation of the FRN by positive and negative RPEs has been suggested as an index of dopaminergic reward processing^17^. As such, the finding that IPEs and RPEs were both reflected in a similar fashion in the FRN provides evidence in favour of the common currency hypothesis^12^, according to which the intrinsic value of information might result from its representation within canonical neural reward-processing circuits. It is important to note that IPEs and RPEs were encoded independently of one another for the task design employed in the present study. A positive RPE—-that is, viewing a black card in the informative stimulus—-could therefore be associated with either a positive, negative, or null IPE depending on the composition of cards preceding and succeeding the event.

Interestingly, our analyses of the trial outcome screens revealed no significant modulation of the FRN by wins versus losses. This is inconsistent both with previous FRN research showing that rewarding outcomes modulated the FRN^21, 23^, and also with the modulation of the FRN by reward prediction errors in response to informative stimuli in the present study. These findings could be due to underpowered statistical analyses, as the task only included one outcome screen event per trial, whereas each trial consisted of several card-elicited reward events, meaning these analyses therefore had comparatively more statistical power. Alternatively, we note that according to one model of information-seeking behaviour, the value of reward-predictive stimuli may exceed the value of the rewarding outcome itself, as a result of the increase in anticipatory utility associated with positive predictive cues^16^.

Several recent findings have challenged the RPE-FRN model of Holroyd and Coles (2002). For instance, Talmi, Atkinson and El-Deredy (2013) found that FRN amplitude, in addition to increasing when reward was unexpectedly withheld, also increased when aversive outcomes were unexpectedly withheld^25^. Since unexpectedly withheld aversion represents a positive RPE, the Holroyd and Coles (2002) model predicts the opposite pattern. Similarly, Hauser and colleagues (2014) reported that the FRN was more strongly associated with the absolute value of RPEs, rather than signed RPEs, and therefore concluded that FRN amplitudes were driven more by surprise than by outcome valence^26^. The results of the present study may suggest an alternative interpretation of these past findings. Our findings demonstrate that the FRN encodes information as well as reward; this finding cannot be explained as a form of surprise encoding, since black and red cards were equally probable for each card draw, and therefore equally surprising according to standard operationalisations of stimulus-bound surprise^27^. Rather, it is possible that past findings demonstrating surprise encoding in the FRN may reflect a complex interaction between RPEs and IPEs. The present study, which averaged across positive and negative IPEs when analysing RPEs, and vice versa when analysing IPEs, had insufficient power to investigate the factorial interaction of RPEs and IPEs. This is an important subject for future research, which could thereby investigate whether there is any asymmetry in IPE encoding as a function of RPE sign, or of RPE encoding as a function of IPE sign.

Under the hypotheses set out above, we did not expect to find any effects of prediction errors on the FRN during non-informative task events. As expected, for these events there was no effect of RPEs on FRN amplitude, as well as no effect of the amount of outcome-relevant information. However, we did find a small but significant difference in amplitude of the FRN elicited by non-informative stimuli during the equivalent of positive and negative IPEs, and this effect was in the same direction as that observed in informative stimuli. One possible explanation for this finding is that, although participants did not receive information about the lottery outcome in the non-informative stimulus, this stimulus may have imparted incidental distributional information. That is, the relative proportions of red and black cards in the non-informative stimulus may have allowed participants to update their beliefs regarding the generative binomial rate of card colours. It is noteworthy in this respect that the effect of IPE in the non-informative stimulus was considerably smaller than the effect in the informative stimulus (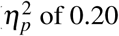 compared to 0.36), and that this non-informative stimulus effect was reduced to a non-significant trend when trials with zero IPEs were included in analysis. These considerations suggest that the non-informative stimulus effect of IPE may have been smaller or less robust than the effect of IPE in the informative stimulus. Alternatively, given the simple and repetitive task design of the present study, another possible explanation for this finding is that participants might also have been unable to suppress tracking the cards ‘as if’ they contained outcome-relevant information. This might have reflected participants’ attempts to assess the generality of the learned associations of card colours. Although we instructed participants that cards in the non-informative stimulus were not predictive of outcomes, participants may nevertheless have attempted to ascertain whether this was truly the case in trials when they chose the non-informative stimulus.

The current study is the first to investigate similarities between information and reward processing in human participants using EEG. Our primary finding, that IPEs and RPEs are both reflected in the FRN, is conceptually consistent with previous studies showing that informative stimuli are encoded in brain regions traditionally associated with reward processing. These include the dopaminergic midbrain and lateral habenula^12, 13^, the orbitofrontal cortex^9^, regions of the striatum^14, 15^ and anterior insula^28^. Under the common currency hypothesis, and premised upon the assumption that the FRN denotes a reward positivity^20^, these findings suggest that acquiring information may be inherently rewarding, regardless of the instrumental use of the information provided. An expected-reward-maximising agent would not give up monetary reward for information that cannot be used to affect task outcomes. However, if information itself has an inherent motivational value, then this value can offset the monetary cost to the participant.

Two distinct neural mechanisms have been proposed which can account for this ‘common currency’ of information and reward. Bromberg-Martin and Hikosaka (2011) posited that the resolution of uncertainty may itself be inherently rewarding, meaning that information has an explicit value unrelated to its instrumental utility for future planning. This explicit value was proposed to manifest in the encoding of IPEs in dopaminergic midbrain neurons. Alternatively, Iigaya and colleagues (2016) noted that animals awaiting the outcome of a lottery might experience anticipatory utility (dread of expected losses and savouring of expected wins). In such a scenario, Iigaya and colleagues (2016) proposed that the absolute value of reward prediction errors elicited by informative stimuli might provide an additive boost to this anticipatory utility. Such a mechanism could result in an apparent encoding of IPEs within canonical reward-processing neurons without the necessity of assuming an explicit value of information, since the card transitions that would be associated with increased/decreased utility in this way are the same as those associated with positive/negative information prediction errors (see Fig. 4C). As such, the findings of the present study are consistent with the mechanisms proposed by both Bromberg-Martin and Hikosaka (2011) and Iigaya et al. (2016). ERP components recorded at the scalp do not measure the activity of dopaminergic midbrain neurons directly, and even direct measures from these brain regions have proven insufficient to distinguish between these two accounts. As such, both models propose viable candidate physiological mechanisms for the observed findings of the present study. More broadly, the role of anticipatory utility in the valuation of information is an important topic for future study. Theories of anticipatory utility can describe individuals’ preferences over deterministic outcomes^29^, sequences of positive outcomes^30^, and two-period decision problems^31^. In determining the relationship of these theories to human information-seeking behaviour, an important topic for future study is the nature of individuals’ preference for non-instrumental information about losses (rather than gains, as in the present study).

In sum, the present study found that human participants exhibited a clear preference for acquiring non-instrumental information about future outcomes. Moreover, the neural encoding of information prediction displayed striking similarities to patterns of encoding of reward prediction errors. An updated decision theory in which information itself is a dimension of stimuli which contributes to their hedonic value-—whether directly, via an explicit valuation of information, or indirectly, via an anticipatory boosting mechanism—-may assist in explaining and predicting patterns of decision making in the presence of reducible uncertainty.

## Methods

### Participants

Participants were 23 healthy, right-handed participants (14 female, 9 male) aged between 18 and 32 years of age (*M* = 23.04, *SD* = 4.15). Participants completed two sessions of a non-instrumental information seeking task (see Fig. 3): a preliminary behavioural session, and an EEG testing session. The present study reports results solely from the EEG testing session; behavioural results from the preliminary session have been previously reported as Experiment 2 in Bennett et al. (2016). Participants received monetary compensation of AUD $10 per session, as well all task winnings up to a maximum of $15 (*M* = $11.48, *SD* = 1.18). All participants provided written informed consent, and research was conducted in accordance with the Declaration of Helsinki. All study protocols were approved by The University of Melbourne Human Research Ethics Committee (ID 1341084).

**Figure 3.**
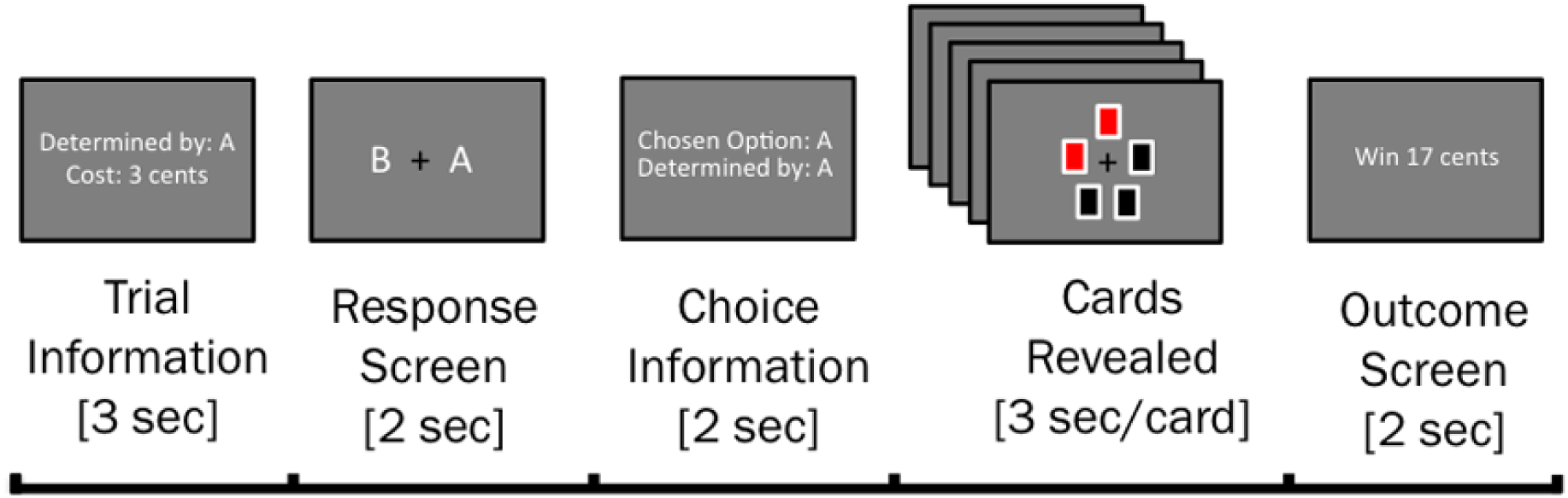
Illustration of trial structure. Participants chose to observe either an informative or a non-informative stimulus concerning the outcome of a monetary lottery. Stimuli were arrays of red and/or black cards. In the informative stimulus, the relative proportion of red and black cards perfectly predicted the lottery outcome (majority black cards: win; majority red cards: loss). In the non-informative stimulus, the relative proportion of card colours was unrelated to the lottery outcome. These options were presented to participants as a choice between Set A and Set B, with the identity of the informative stimulus pseudo-randomised across trials. Participants were also informed of the cost of observing the informative stimulus. After the participant’s choice, cards were drawn one-by-one at a constant rate. Once all cards were drawn, the outcome of the monetary lottery was revealed to participants.

### Protocol and apparatus

Before commencing the preliminary behavioural session, participants received verbal and written instruction of the task and were permitted to complete a practice task. Participants were informed that their choice of stimulus on each trial would not affect the likelihood of winning the lottery, and that the probability of winning and losing was equal on each trial.

The task was presented on a Dell P2210 LCD monitor (1680 x 1050 screen resolution; refresh rate 60 Hz) using the Psychophysics Toolbox^32^. On each trial, the participant chose between an informative and a non-informative stimulus by using the index finger of their right hand to press either the left or the right button of a five-button Cedrus response box. The left-right response mapping was pseudo-randomised across trials. Participants completed 7 blocks of 16 trials each while EEG was recorded, with a total testing duration of approximately 50 minutes.

In order to ensure that participants maintained attention on the chosen stimulus as it was revealed, a small number of trials (approx. 10%) were ‘catch trials’, in which one card in the chosen stimulus was drawn to reveal a white X rather than a red or black card. Participants were instructed to respond to this attention check by pressing any button within 1.5 seconds. Failure to do so resulted in the deduction of $1 from overall winnings. This ensured that participants did not disengage from the task during stimulus presentation, and attended equally to both stimulus types.

In line with previous research using this task^6^, it was determined a priori that participants who failed to respond to more than two catch trials would be deemed to have failed an attention check. One participant failed to respond on four catch trials, and was therefore excluded from all further analysis. The remaining participants showed good levels of task engagement as measured by successful responses to catch trials (M = 98.11%, SD = 3.57%).

### EEG data acquisition and preprocessing

EEG data were acquired from 64 Ag/AgCl active scalp electrodes located according to the International 10-20 system. Data were recorded at a sampling rate of 512 Hz using a BioSemi ActiveTwo system using an implicit reference during recording, and were linearly detrended and re-referenced offline to an average of left and right mastoids. The electrooculogram was recorded from two infraorbital electrodes horizontally adjacent to and below the left eye.

EEG data were preprocessed using EEGLAB^33^, according to a semi-automated preprocessing pipeline^34, 35^. Data were high-and low-pass filtered at 0.1 Hz and 70 Hz respectively, and notch filtered from 45-55 Hz to remove background electrical noise. Data were segmented into epochs from 1000ms before to 1000ms after events of interest, and baseline corrected using a 100ms pre-stimulus baseline. An Independent Component Analysis (ICA) as implemented in EEGLAB was used to identify and remove components of the data related to eyeblink and saccade artefacts. Noisy data channels were interpolated using a spline interpolation routine; no interpolated data channels were included in final ERP analyses. Finally, an impartial artefact screening procedure automatically excluded all epochs in which maximum/minimum amplitudes exceeded 200 mV.

### EEG data analysis

ERP analyses were conducted using ERPLAB^36^. In line with previous research, FRN amplitudes were calculated for each condition as the mean amplitude from 200 to 350 milliseconds post-stimulus at the five fronto-central channels Fpz, AFz, Fz, FCz, and Cz^23^. These five channels are located above the medial frontal cortex, a candidate generator for the FRN^17, 21^. For all analyses, ANOVA degrees of freedom were adjusted using the Greenhouse-Geisser correction where the assumption of sphericity was violated.

Finally, since behavioural results showed large differences in strategies between individuals, the number of epochs available for different ERP analyses differed between participants. ERP analyses therefore only included data from participants who had at least 20 epochs of each event type under consideration^19^. See Table S1 in the Supplementary Materials for a summary of the number of participants and trials included in each analysis.

### Quantification of computational variables

Epochs were binned for ERP analysis according to three different computational variables: positive/negative RPEs, positive/negative IPEs, and positive/negative information. Each of these variables was calculated according to the difference between win probabilities before and after observation of a card. This quantity was calculated based on the binomial probability that black cards would be in the majority in the informative stimulus, given conditional independence of successive card draws:

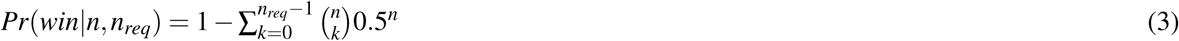

Where *n* is the number of cards remaining to be drawn, and *n*_*req*_ is the number of additional black cards required for a majority given *n*_*black*_ already drawn:

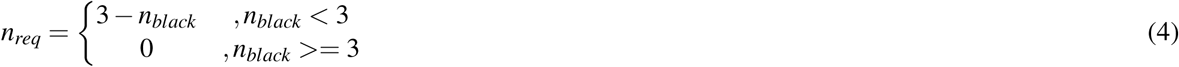

By definition, no information is imparted by cards in the non-informative stimulus, and in this case was always equal to 0.5.

The reward prediction error associated with each card was calculated as the difference between the participant’s expected lottery winnings prior to and following the card draw (see Equation 1).

Following Shannon (1948)^37^, the information content of a stimulus was defined as the entropy difference between posterior and prior beliefs:

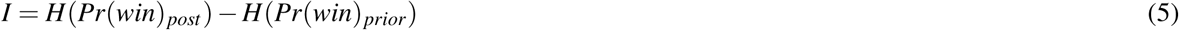

With entropy defined as the binary entropy function:

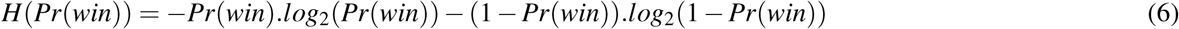

Given *n* and *n*_*req*_, the expected information content of any card prior to its being revealed can therefore be calculated as the average of the amount of information that would be associated with one additional red card and the amount of information that would be associated with one additional black card:

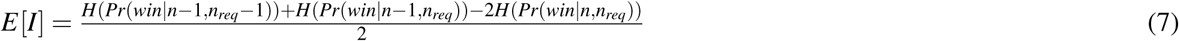

This allows for IPEs to be calculated as per Equation 2. See Fig 4 for a schematic overview of card transitions indicating events associated with positive versus negative RPEs and IPEs, and the differences between the two. Positive/negative RPEs indicate an increase in the likelihood of winning/losing the lottery, respectively, whereas positive/negative IPEs indicate that an event conveyed more/less information about the outcome than expected, regardless of whether that outcome was a win or a loss. Given that there was an equal probability of observing a red versus a black card at every point in time (i.e. each state transition in Fig. 4 was equiprobable), positive and negative RPEs and IPEs were therefore independent of one another for the task design used in the present study.

**Figure 4.**
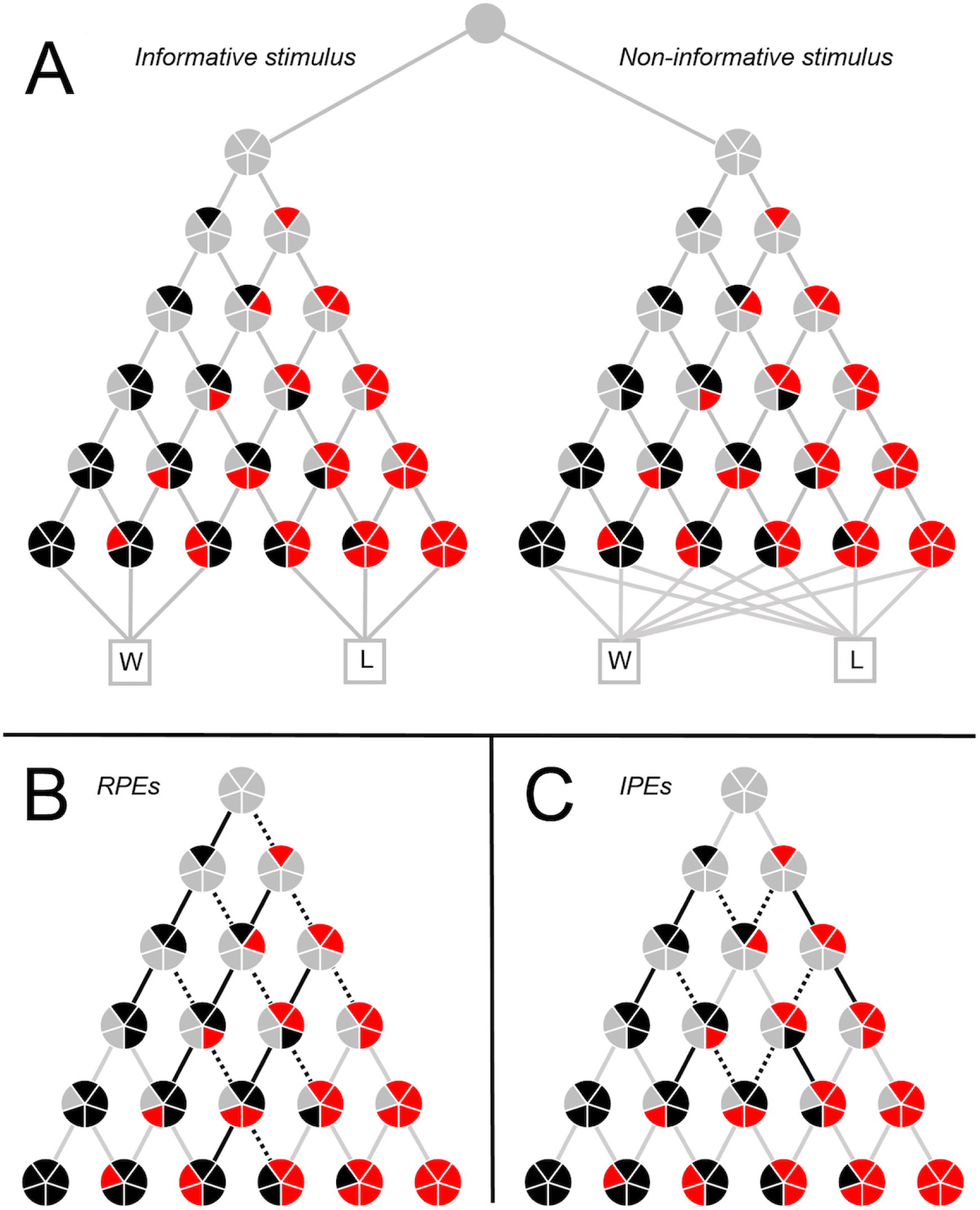
Schematic overview of task events. Segment colours within each circle denote the number of black and red cards visible at any point in time, whereas lines denote transitions between states (card draws). (A) Overall schematic including both informative (left) and non-informative (right) stimuli. Stimuli were perceptually identical, and differed only in that the majority colour in the informative stimulus perfectly predicted lottery outcome (win, W; loss, L). (B) Informative stimulus schematic, with positive reward prediction errors (RPEs) denoted by solid lines, negative RPEs by dashed lines, and zero RPEs by grey lines. (C) Informative stimulus schematic, with positive information prediction errors (IPEs) denoted by solid lines, negative IPEs by dashed lines, and zero IPEs by grey lines.

## Data availability

Data is available upon reasonable request from the corresponding author.

## Acknowledgements

The authors wish to thank Hayley Warren and William Turner for help with data acquisition and feedback on the manuscript. This work was supported by a Faculty of Business and Economics (University of Melbourne) Strategic Initiatives Grant 2011 to S.B. and C.M and an Australian Research Council (ARC) Discovery Early Career Researcher Award (DECRA) to S.B. (DE140100350).

## Author contributions statement

D.B., C.M. and S.B. designed the task. M.B. and D.B. collected the data. M.B. and D.B. analysed the data. M.B., D.B., C.M. and S.B. wrote the paper.

## Additional information

## References

1. Kidd, C. & Hayden, B. Y. The psychology and neuroscience of curiosity. Neuron 88, 449–460 (2015).

2. Raiffa, H. & Schlaifer. Applied Statistical Decision Theory (Division of Research, Harvard Business School, Boston, 1961).

3. Howard, R. A. Information value theory. IEEE Transactions on Syst. Sci. Cybern. 2, 22–26 (1966).

4. Lawrence, D. B. The Economic Value of Information (Springer Science & Business Media, 2012).

5. Lanzetta, J. T. & Driscoll, J. M. Preference for information about an uncertain but unavoidable outcome. J. Pers. Soc. Psychol. 3, 96 (1966).

6. Bennett, D., Bode, S., Brydevall, M., Warren, H. & Murawski, C. Intrinsic valuation of information in decision making under uncertainty. PLoS Comput. Biol. 12, e1005020 (2016).

7. Zhu, J.-Q., Xiang, W. & Ludvig, E. A. Information seeking as chasing anticipated prediction errors. In Proceedings of the 39th Annual Meeting of the Cognitive Science Society (2017).

8. Zentall, T. R. & Stagner, J. Maladaptive choice behaviour by pigeons: an animal analogue and possible mechanism for gambling (sub-optimal human decision-making behaviour). Proc. Royal Soc. Lond. B: Biol. Sci. 278, 1203–1208 (2011).

9. Blanchard, T. C., Hayden, B. Y. & Bromberg-Martin, E. S. Orbitofrontal cortex uses distinct codes for different choice attributes in decisions motivated by curiosity. Neuron 85, 602–614 (2015).

10. Vasconcelos, M., Monteiro, T. & Kacelnik, A. Irrational choice and the value of information. Sci. Reports 5, srep13874 (2015).

11. Grant, S., Kajii, A. & Polak, B. Intrinsic preference for information. J. Econ. Theory 83, 233–259 (1998).

12. Bromberg-Martin, E. S. & Hikosaka, O. Lateral habenula neurons signal errors in the prediction of reward information. Nat. Neurosci. 14, 1209–1216 (2011).

13. Bromberg-Martin, E. S. & Hikosaka, O. Midbrain dopamine neurons signal preference for advance information about upcoming rewards. Neuron 63, 119–126 (2009).

14. Kang, M. J. et al. The wick in the candle of learning: Epistemic curiosity activates reward circuitry and enhances memory. Psychol. Sci. 20, 963–973 (2009).

15. Jepma, M., Verdonschot, R. G., Van Steenbergen, H., Rombouts, S. A. & Nieuwenhuis, S. Neural mechanisms underlying the induction and relief of perceptual curiosity. Front. Behav. Neurosci. 6 (2012).

16. Iigaya, K., Story, G. W., Kurth-Nelson, Z., Dolan, R. J. & Dayan, P. The modulation of savouring by prediction error and its effects on choice. eLife 5, e13747 (2016).

17. Holroyd, C. B. & Coles, M. G. The neural basis of human error processing: reinforcement learning, dopamine, and the error-related negativity. Psychol. Rev. 109, 679 (2002).

18. Cohen, M. X. Neurocomputational mechanisms of reinforcement-guided learning in humans: a review. Cogn. Affect. & Behav. Neurosci. 8, 113–125 (2008).

19. Hajcak, G., Moser, J. S., Holroyd, C. B. & Simons, R. F. It’s worse than you thought: The feedback negativity and violations of reward prediction in gambling tasks. Psychophysiol. 44, 905–912 (2007).

20. Holroyd, C. B., Krigolson, O. E. & Lee, S. Reward positivity elicited by predictive cues. Neuroreport 22, 249–252 (2011).

21. Cohen, M. X., Elger, C. E. & Ranganath, C. Reward expectation modulates feedback-related negativity and eeg spectra. Neuroimage 35, 968–978 (2007).

22. Miltner, W. H., Braun, C. H. & Coles, M. G. Event-related brain potentials following incorrect feedback in a time-estimation task: Evidence for a “generic” neural system for error detection. J. Cogn. Neurosci. 9, 788–798 (1997).

23. Hajcak, G., Moser, J. S., Holroyd, C. B. & Simons, R. F. The feedback-related negativity reflects the binary evaluation of good versus bad outcomes. Biol. Psychol. 71, 148–154 (2006).

24. Walsh, M. M. & Anderson, J. R. Learning from experience: event-related potential correlates of reward processing, neural adaptation, and behavioral choice. Neurosci. & Biobehav. Rev. 36, 1870–1884 (2012).

25. Talmi, D., Atkinson, R. & El-Deredy, W. The feedback-related negativity signals salience prediction errors, not reward prediction errors. J. Neurosci. 33, 8264–8269 (2013).

26. Hauser, T. U. et al. The feedback-related negativity revisited: new insights into the localization, meaning and network organization. Neuroimage 84, 159–168 (2014).

27. Mars, R. B. et al. Trial-by-trial fluctuations in the event-related electroencephalogram reflect dynamic changes in the degree of surprise. J. Neurosci. 28, 12539–12545 (2008).

28. Preuschoff, K., Quartz, S. R. & Bossaerts, P. Human insula activation reflects risk prediction errors as well as risk. J. Neurosci. 28, 2745–2752 (2008).

29. Loewenstein, G. Anticipation and the valuation of delayed consumption. The Econ. J. 97, 666–684 (1987).

30. Loewenstein, G. F. & Prelec, D. Preferences for sequences of outcomes. Psychol. Rev. 100, 91 (1993).

31. Caplin, A. & Leahy, J. Psychological expected utility theory and anticipatory feelings. Q. J. Econ. 116, 55–79 (2001).

32. Brainard, D. H. & Vision, S. The psychophysics toolbox. Spatial Vis. 10, 433–436 (1997).

33. Delorme, A. & Makeig, S. Eeglab: an open source toolbox for analysis of single-trial eeg dynamics including independent component analysis. J. Neurosci. Methods 134, 9–21 (2004).

34. Bennett, D., Murawski, C. & Bode, S. Single-trial event-related potential correlates of belief updating. eNeuro 2, ENEURO–0076 (2015).

35. Bode, S., Bennett, D., Stahl, J. & Murawski, C. Distributed patterns of event-related potentials predict subsequent ratings of abstract stimulus attributes. PloS One 9, e109070 (2014).

36. Lopez-Calderon, J. & Luck, S. J. Erplab: an open-source toolbox for the analysis of event-related potentials. Front. Hum. Neurosci. 8 (2014).

37. Shannon, C. A mathematical theory of communication. Bell Syst. Tech. J. 843, 379–423 and 623–656 (1948).

